# Trapping DNA-radicals with DMPO reduced hypochlorous acid-induced 8-oxo-7,8-dihydro-2’-deoxyguanosine and mutagenesis in lung epithelial cells

**DOI:** 10.1101/2024.06.18.599657

**Authors:** D.C. Ramirez, S.E. Gomez Mejiba

## Abstract

Irritation causes the recruitment and activation of neutrophils in the stressed airways. This process is known as neutrophilic inflammation. This process results in myeloperoxidase (MPO), an enzyme contained inside neutrophil azurophilic granules, being released as neutrophil extracellular traps (NETs), which also contain genomic DNA, modified histones, and other proteins. In the airways, released MPO can be taken up by bystander tissue epithelial cells. MPO is the only mammalian peroxidase enzyme that under physiological conditions produces hypochlorite (HOCl). Intracellularly produced HOCl may damage the cell genome, with the intermediacy of DNA-centered free radicals, which upon reaction with molecular oxygen decay to mutagenic end-oxidation products, such as 8-oxo-7,8-dihydro-2’ –deoxyguanosine (8-oxo-dGuo). Herein, we aimed to test whether HOCl-induced DNA-centered radicals precede the oxidation of DNA and mutagenesis in A549 human lung epithelial cells as an *in vitro* model that resembles neutrophilic inflammation in irritated airways. Interestingly, by trapping HOCl-induced DNA-centered radicals, the nitrone spin trap 5,5-dimethyl-1-pyrroline *N*-oxide (DMPO) blocks the formation of 8-oxo-dGuo and possibly other end-oxidation products, forming DNA-DMPO nitrone adducts, thus reducing mutagenesis in the hypoxanthine phosphoribosyl transferase (*hrpt)* gene, one of the most sensitive genes to oxidative damage. P53 is a transcription factor known as the master regulator of the cell response to genomic damage. By trapping DNA-centered radicals, DMPO also blocks the translocation of p53 to the cell nucleus, suggesting that by trapping DNA-centered radicals with DMPO, end-oxidation products are prevented, and the cell response to genomic damage is not sensed. DMPO traps DNA-centered radicals, reduces 8-oxo-dGuo accumulation, and blocks *hrpt* gene mutation. Trapping DNA-centered radicals to reduce the accumulation of HOCl-induced mutagenic end-oxidation products in the genome of bystander cells, which have taken MPO from the inflammatory milieu, will provide new therapeutic avenues to reduce genotoxic damage at sites of neutrophilic inflammation, such as in the irritated airways.

## INTRODUCTION

Neutrophilic inflammation is a normal acute tissue response to irritation. This irritation can be caused by biological, physical, mechanical, or metabolic cues [1]. The cell damage response to irritation results in the release of chemokines by tissues and resident macrophages that attract neutrophils from blood into the irritated tissue following a chemokine gradient [2]. At sites of irritation, recruited neutrophils are activated with the production of superoxide radicals for activation of NADPH oxidase [3] and fusion of azurophil granules containing myeloperoxidase (MPO, donor hydrogen peroxide, oxidoreductase, E.C. 1.11.1.7) with other components, including elastase, histones and genomic DNA, forming neutrophil-extracellular traps (NETs) [4]. In the extracellular milieu, DNAses release NET-associated MPO that can be taken up by surrounding tissue cells [5, 6]. Inside cells, MPO is still active where it can produce large amounts of HOCl [7, 8], a powerful and random oxidant that can oxidize any macromolecule depending on proximity to its source (discussed in [9]).

Recruitment and activation of neutrophils are named neutrophilic inflammation [1], which is, from a biochemical standpoint, different from neutrophilic infiltration, which is a term reserved for a histological pattern [10]. Neutrophilic inflammation can result in mutations leading to cell transformation and carcinogenesis by HOCl-driven oxidatively generated modifications to genomic DNA [11]. Previously, we found that a bolus addition of HOCl or HOCl generated by the biochemical system MPO/H_2_O_2_/Cl^−^ can oxidize calf-thymus DNA with the formation of DNA-centered radicals. Furthermore, inside cells loaded with MPO and in HL-60 cells that overexpress MPO, DNA radicalization can be determined using the cell-permeable nitrone spin trap 5,5-dimethyl-1-pyrroline *N*-oxide (DMPO), which forms DNA-DMPO radical adducts that can be measured using immunospin trapping (IST) technology [8]. A detailed protocol for measuring DNA radicalization using IST has been published [12], and IST technology is reviewed in [8]. Interestingly, DNA radicalization precedes 8-oxo-7,8-dihydro-2’ – deoxyguanosine (8-oxo-dGuo) during hydroxyl radical and carbonate radical anion reactions with genomic DNA [13]. 8-oxo-dGuo is a molecular marker of free radical-induced DNA damage and carcinogenesis [14].

Neutrophilic inflammation, in particular the activity of MPO, is thought to cause genomic damage in tissues and may be involved in cell transformation [15–18]. Once MPO is taken up by surrounding tissue cells [5] and at sites of inflammation where the concentration of H_2_O_2_—a nonfree radical cell-permeable reactive oxygen species, is produced in the range of 30-200 μM, intracellular production of HOCl is possible [16]. Indeed, in HL-60 cells, MPO is present in the cell nuclei, whereas in A549 epithelial cells, it accumulates close to the nuclear envelope, and intracellular concentrations of Cl^−^ are between 5-100 mM [7]. HOCl reacts with nuclear DNA or DNA-forming NETs to produce DNA chloramines that decay, forming DNA-(*N*)-centered radicals [19, 20]. The unpaired electron can be delocalized to other atoms in nucleotides, resulting in a wide range of oxidation and chlorination products (bases and sugar) [21]. *In vitro*, DNA radicals and those formed when nucleotides, nucleosides, and bases are reacted with HOCl have been previously observed using electron-spin resonance [20]. The rate constant for the reaction of nucleotides and their components with HOCl has been reported [22].

DNA damage is sensed by the transcription factor p53–the master regulator of cell response to DNA damage, which is accumulated in the nucleus where it binds to response elements and induces the expression of genes involved in arresting the cell cycle until repair mechanisms are triggered. This response includes the expression of p21 and inhibition of cyclin/CDK complex formation [23, 24]. The *hypoxanthine phosphoribosyl transferase* (*hrpt*) gene is one of the most sensitive genes to oxidative DNA damage[25]. The *hrpt* gene is mapped on the X chromosome of mammalian cells, and it is used as a model gene to investigate gene mutations in mammalian cell lines [26]. In addition, there is no doubt that neutrophilic inflammation may be involved in causing mutations and further cell transformation and carcinogenesis [11], for example, in the lung exposed to environmental stressors such as insect products, endotoxin, and particles [17, 27] and SARS-CoV-2 infection in COVID-19 patients’ airways [28], but the mechanism connecting DNA radical formation and mutations remains to be fully elucidated.

Herein, we show that HOCl produced by MPO inside A549 cells causes, sequentially, DNA radicalization, 8-oxo-dGuo formation, nuclear accumulation of p53, and mutations of the *hrpt* gene. After DNA radicalization, the nitrone spin trap DMPO blocks all these processes, thus preventing the mutation of the *hrpt* gene.

## METHODS

### Treatment with purified human myeloperoxidase

*Preparation of MPO*—MPO was prepared as we previously described in [7]. Briefly, 100 μg of highly purified human MPO (Athens Research and Technology, cat# 16-14-130000) was dissolved in 200 μL of ultrapure water and dialyzed overnight against 2 L of 10 mM Chelex-sodium phosphate buffer, pH 7.4. To produce inactivated MPO, the enzyme was incubated with 100 μM KCN or 500 μM 3-amino-1,2,4-triazole (ATZ, Sigma, Cat# A-8056) for 30 minutes at 37 °C. Before the addition of MPO (native or inactivated) to the cell culture medium, the solution was sterilized by passing it through a 0.22 μm nylon syringe filter. The UV‒visible spectrum was used to determine the concentration, identity, and purity of the MPO. The Soret band with a peak maximum at 430 nm (178,000 M^-1^ cm^-1^ for the MPO homodimer at pH 7.4) is characteristic of the specific heme-prosthetic group in MPO. The Rheinheitzahl value (*A*_430_/*A*_280_ ratio) from the UV‒visible spectra provided an estimate of the purity of MPO relative to total protein. The preparations of native MPO used in our experiments had a Rheinheitzahl value of 0.8. MPO inactivation was measured by the oxidation of guaiacol to tetraguaiacol at 470 nm (ɛ_470 nm_ = 26.6 mM^−1^ cm^−1^) [7]. To adjust MPO concentrations, we performed protein measurements by using the BCA protein assay.

### The biochemical systems for the generation of DNA-centered radicals

To determine the generation of DNA-nitrone adducts in a biochemical system, we incubated calf thymus double-stranded deoxyribonucleic acid (dsDNA, ∼10,000–15,000 kDa) as its sodium salt (cat# D3664) with 5 μM MPO in 10 mM sodium phosphate buffer containing 140 mM Cl^−^, as NaCl, pH 7.4. The generation of HOCl was started by a bolus addition of 50 μM H_2_O_2_ (ɛ_240_ =

43.6 M^−1^ cm^−1^). Fifteen minutes later, 50 μM DMPO (Dojindo Molecular Technologies Inc., Japan, Cat# D04810, ɛ_228_ = 7,800 M^−1^ cm^−1^) was added to trap DNA-centered radicals resulting from DNA-chloramine decay [20]. The total reaction volume was 100 μL. After 1 h of incubation, the reaction was stopped by adding 1 mM methionine. MPO inhibitors (KCN, 4-aminobenzoic acid hydrazide (ABAH, Cayman Chemicals, cat# 4845) or salicyl hydroxamic acid (SHA, Sigma cat# S607)) or scavengers of HOCl (methionine or taurine) were added 15 minutes before H_2_O_2_ addition.

### A549 cell culture and loading with MPO

The lung type-2 epithelial cell line A549 (ATCC, Cat# CCL-185^TM^) was cultured in Ham’s F12 medium (Lonza Biosciences, Cat# 12615F) containing 10% heat-inactivated fetal calf serum (FCS, Gibco, Ca#A3840101). Depending on the experiment, A549 cells were cultured in 96-well or 6-well plates with a 5 mm-round cover slide in their bottom. Cells were grown to 85% confluence before each experiment. A549 epithelial cells were loaded with MPO by incubation in fresh medium with 5% FCS and 10 nM native MPO or inactivated MPO. After 24 h of incubation, the cells were rinsed with HBSS^−^ (HBSS without Ca^2+^ or Mg^2+^, Lonza, cat# 04-315Q) and incubated in HBSS^+^ with 50 μM H_2_O_2_ or HBBS^+^ containing 5 mM glucose, and HOCl generation started with 0.5 mIU/mL glucose oxidase (GO) for 15 min, followed by the addition or not of 50 mM DMPO.

### Neutrophil isolation and coculture with A549 lung epithelial cells

Isolation and coculture of human neutrophils with A549 epithelial cells were performed as described in [7]. Briefly, confluent monolayers of A549 cells and 10^6^ isolated human neutrophils in coculture were incubated in 2 ml of HBSS^+^ on round 5 mm cover slides in 6-well tissue culture plates at 37 °C. To generate NETs, we treated the coculture with 100 ng/ml phorbol 12-myristate 13-acetate (PMA, Sigma, cat# P8139) for 1 h in the incubator. Finally, the monolayers were rinsed 4 times with HBSS^−^ before staining and confocal microscopy analyses.

### Confocal microscopy

For laser confocal microscopy, cells were cultured on 5 mm glass-cover slides in 6-well plates. Immuno-staining of MPO, p53, and histone H2B was performed following a procedure similar to that followed in our previous study [7]. Briefly, after treatments, the slide was rinsed and fixed with 4% paraformaldehyde for 15 min at 37 °C and then permeabilized with 0.2% Triton X-100 at room temperature for 5 min, followed by blocking with the Image-iT^TM^ FX supersignal enhancer (Thermo Fisher Scientific, cat# I36933). Fixed cells were incubated overnight at 4 °C with the primary antibodies, washed, and then incubated with the secondary antibodies at 37 °C for 1 hour. The primary antibodies used were anti-human p53 (Epitomics, cat# 1047-1, rabbit monoclonal anti-human p53, C-term, dilution 1:100), rabbit polyclonal anti-human MPO (Athens Research and Technology, cat# 01-14-130000, dilution, 1:500), and mouse monoclonal anti-human histone H2B (Abcam, clone 4G6, cat# H00008349-M06, dilution 1:100). The secondary antibodies used were goat anti-rabbit (Fc) conjugated to Alexa Fluor 488 or sheep anti-mouse IgG (Fc) conjugated to Texas Red. Finally, slides were washed with PBS and mounted using Prolong Gold anti-fade mountant with 4,6-diamino-2-phenylindole (Thermo Fisher Scientific, cat# P36931), and the preparation was examined with a Leica SP2 MP Confocal Microscope with a 63 x 1.4 oil immersion objective. Single plane or z-stack images were acquired and analyzed using LSM 5 image examiner software.

### Determination of H_2_O_2_-triggered HOCl production inside MPO-loaded A549 epithelial cells

Intracellularly generated HOCl was determined using luminol as a luminescent probe [29]. After loading the cells with MPO in a 96-well clear-bottom black microplate, monolayers were rinsed, and 50 μL of 10 μM luminol (Sigma, cat# 123072) was added to 100 mM Chelex-sodium phosphate buffer, pH 7.4. When resveratrol (Sigma, cat# R5010) or taurine (Sigma, cat# T0625) was tested, they were added together with the luminol reagent. The intracellular generation of HOCl was started by adding 100 μL of 100 mM sodium phosphate buffer with 100 μM H_2_O_2_, and the luminescence was read in a microplate reader within 5 min of mixing the reagents at 15–22 °C. When H_2_O_2_ was generated by the G/GO system, 1 mIU/mL GO (Sigma, cat# G7141) was added to the medium (HBSS^+^ with 5.6 mM glucose), and incubation was continued in the plate reader for 5 minutes at 37 °C, followed by luminescence measurements. After reading the luminescence, dsDNA bound to the bottom of the plate was measured. Similar data were found using the fluorescent probe 3’-(p-aminophenyl) fluorescein (APF, Invitrogen, cat# A36003), which is a fairly specific probe for HOCl.

### Quantification of double-stranded DNA bound to microtiter plates

To quantify dsDNA bound to black microplates, we used 4,6-diamino-2-phenylindole (DAPI, Thermo Fisher Scientific, cat# D1306), which intercalates DNA and fluoresces (λ_ex_= 345 nm/λ_em_ = 458 nm) as described in [7].

### Determination of cell viability and GAPDH activity

Cell viability was determined using the 3-(4,5-dimethylthiazol-2-yl)-2,5-diphenyltetrazolium bromide tetrazolium reduction assay (MTT) [30] and lactic dehydrogenase release assays. Glyceraldehyde-3-phosphate dehydrogenase (GAPDH) is a critical enzyme for cell survival and is extremely sensitive to HOCl-induced oxidative damage [9]. Thus, our experiments were complemented by the determination of GAPDH activity in homogenates to ensure early damage to energy metabolism.

### Extraction of DNA and determination of 8-oxo-dGuo and DNA-DMPO nitrone adducts

After treatments, the cells were rinsed with HSSB^-^, and genomic DNA was isolated using the protocol we previously published [12]. The contents of 8-oxo-dGuo and DMPO-DNA nitrone adducts were determined by an enzyme-linked sorbent immunoassay (ELISA) as we have previously described in [13]. These technologies were recently reviewed in [8].

### 6-Thioguanine mutagenesis assay

The 6-TG mutagenesis assay can detect a wide range of chemicals capable of causing DNA damage that leads to gene mutation [26]. The 6-TG mutagenesis assay is based on the fact that mutations that destroy the functionality of the *hrpt* gene and/or protein are detected by positive selection using a toxic purine analog (6-thioguanine, 6-TG). After 14 days of incubation with 10 μg/mL 6-TG (Sigma, cat# A4882), only mutant cells survived and proliferated, and they could be measured using the trypan blue exclusion assay. A549 cells loaded with active MPO and treated or not with the G/GO system and in the presence or absence of DMPO were screened for 6-TG resistance by using the trypan blue exclusion assay. See **Fig. 6A** for a scheme of the workflow used for the *hrpt* mutagenesis assay.

### Statistics

Relative light units (RLU) or RLU/μg of dsDNA are reported as the mean values ±s.e.m. Differences between pairs were determined by Student’s *t* test and between treatments and control by one-way analysis of variance with Dunnett’s post hoc testing. Differences were considered statistically significant at *p* < 0.05.

## RESULTS

To show HOCl-induced radicalization of DNA, we incubated calf thymus DNA with MPO and H_2_O_2_ for 15 minutes followed by the addition of 50 mM DMPO to trap DNA-centered radicals resulting from DNA-chloramine decay (**Fig. 1A**). Once formed, DNA-centered radicals can be scavenged by reduced glutathione (GSH) or L-ascorbate [7]. We measured DNA-DMPO nitrone adducts by ELISA (**Fig. 1B**). The addition of KCN or inhibitors of MPO, such as ABAH or SHA, resulted in a reduction in the DNA-DMPO nitrone adduct content. Taurine and methionine reduce DNA radical formation by reacting with HOCl before it can react with DNA.

**Figure 1:**
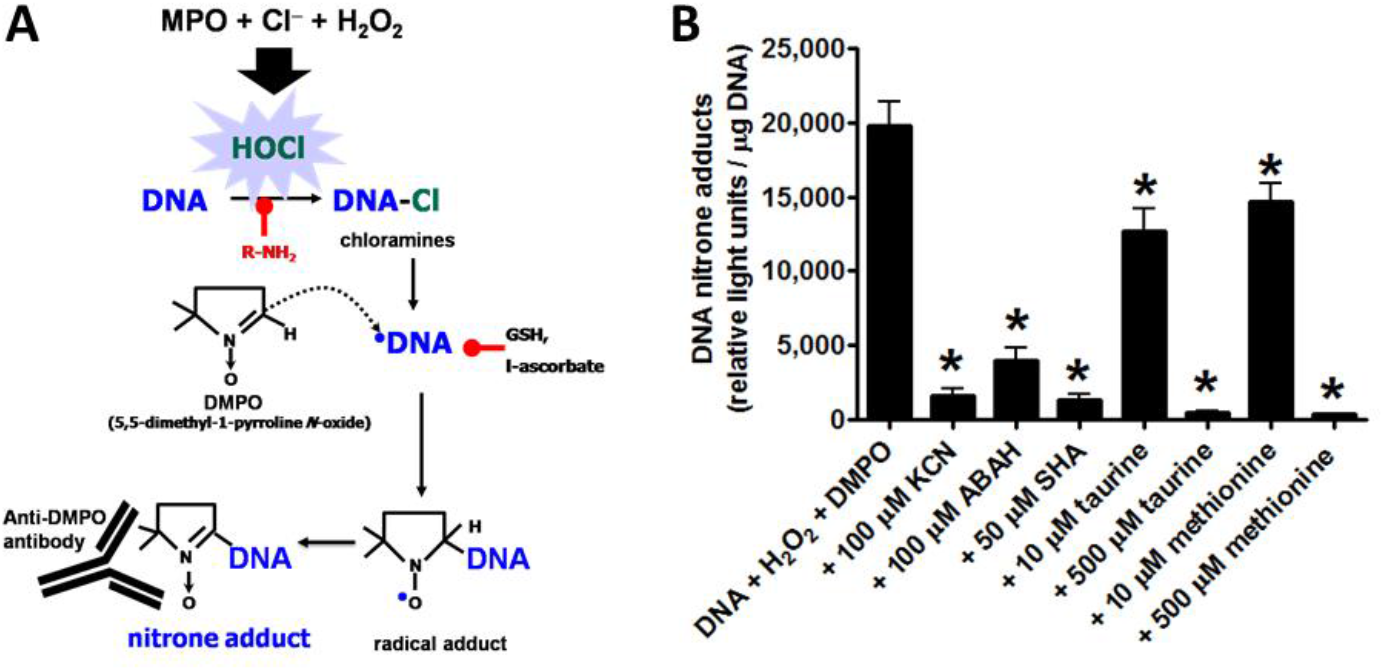
Calf-thymus DNA radicalization by the MPO/H_2_O_2_/Cl^−^ biochemical system and its measurement using immunospin trapping. **A**) Schematic pathway to produce DNA radicalization by HOCl generated by the MPO/H_2_O_2_/Cl system. HOCl reacts with DNA to produce transient DNA chloramines that quickly decay, forming DNA-*N*-centered radicals. These radicals are trapped by DMPO-forming DNA-DMPO radical adducts that decay, forming DMPO-nitrone adducts (see [8] for a recent review). These are recognized by the anti-DMPO antibody. In cells, GSH and L-ascorbate can compete with DMPO for DNA-centered radicals. **B**) ELISA for DNA-DMPO nitrone adducts formed when calf-thymus DNA was incubated with MPO in a buffer containing physiological concentrations of Cl^−^ and HOCl formation was started by H_2_O_2_. Potassium cyanide (KCN), 4-aminobenzoic acid hydrazide (ABAH), salicylhydroxamic acid (SHA), taurine, or methionine was added before HOCl formation with 50 μM H_2_O_2_. Data are shown as the mean values ± s.e.m. from three separate experiments in triplicate. Asterisks indicate p<0.05.

To simulate what occurs at sites of neutrophilic inflammation in the lung, we incubated A549 lung epithelial cells with PMA-activated neutrophils (**Fig. 2A**). Neutrophil activation is observed as the formation of NETs in which MPO and other proteins, along with other nuclear components (histones and genomic DNA), are released, forming NET-like structures and accumulation of MPO close to the nuclear membrane of A549 cells (**Fig. 2B**). Z-stacks of images allowed the observation of NETs and nuclear accumulation of MPO inside A549 cells.

**Figure 2:**
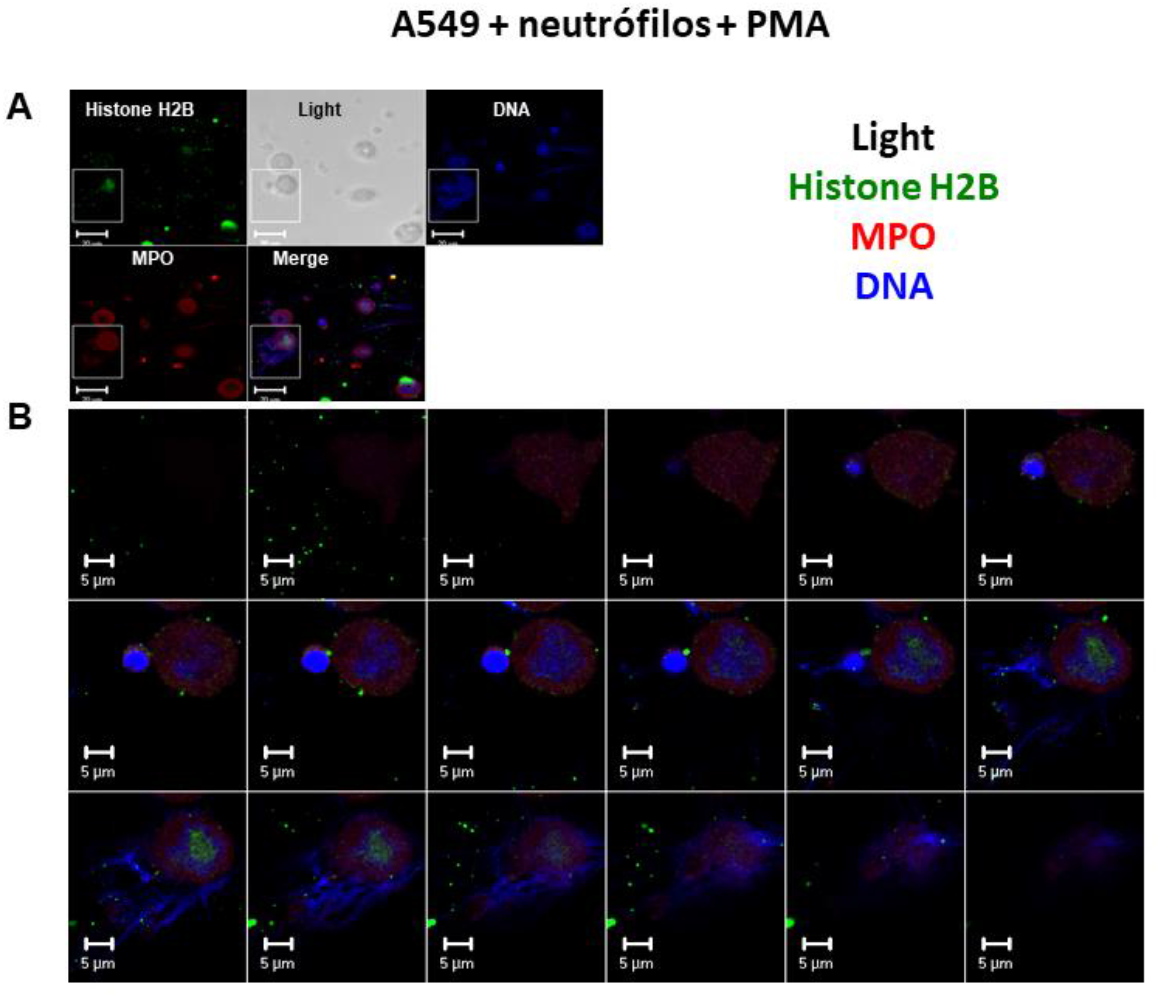
Coculture of A549 lung epithelial cells with PMA-activated neutrophils showing NETs in sequential planar images. **A**) Scanning confocal microscopy image showing the localization of histone H2B, MPO, and genomic DNA. Measurement bars are 20 μm. The square shows a merged image of the interaction of a NET generated during neutrophil activation with one A549 cell. **B**) Eighteen z-stack of planar images of the image shown in the square of panel A. Measurement bars are 5 μm. Images are representative of three individual experiments.

To simulate neutrophilic inflammation and accumulation of MPO inside cells, we incubated A549 human lung epithelial cells with highly purified native human MPO and imaged its cell localization by using confocal microscopy imaging (**Fig. 3A**). This is a good model to study HOCl-induced DNA radicalization in cells because any protein added to cell culture is taken up by cells. Then, we measured the highest concentration of H_2_O_2_ that does not cause toxicity in A549 cells loaded with MPO (**Fig. 3B**) after incubation for 15 minutes. A bolus addition of 50 μM H_2_O_2_ did not produce changes in the cell viability of MPO-loaded A549 cells as assessed by the MTT assay. LDH release and neutral red assays gave similar results (data not shown). In any case, intracellularly generated HOCl can oxidize any protein depending on proximity, including GAPDH [9]—a highly sensitive enzyme critically involved in the glycolytic metabolism of glucose. A dose of 50 μM H_2_O_2_ caused a significant loss of enzyme activity, suggesting the beginning of damage to cell metabolism (**Fig. 3B**). A concentration of 100 μM H_2_O_2_ added to the culture of A549 epithelial cells, without MPO, did not affect either GAPDH activity or cell viability (data not shown), suggesting that the observed effects are caused by HOCl produced inside the cell.

**Figure 3:**
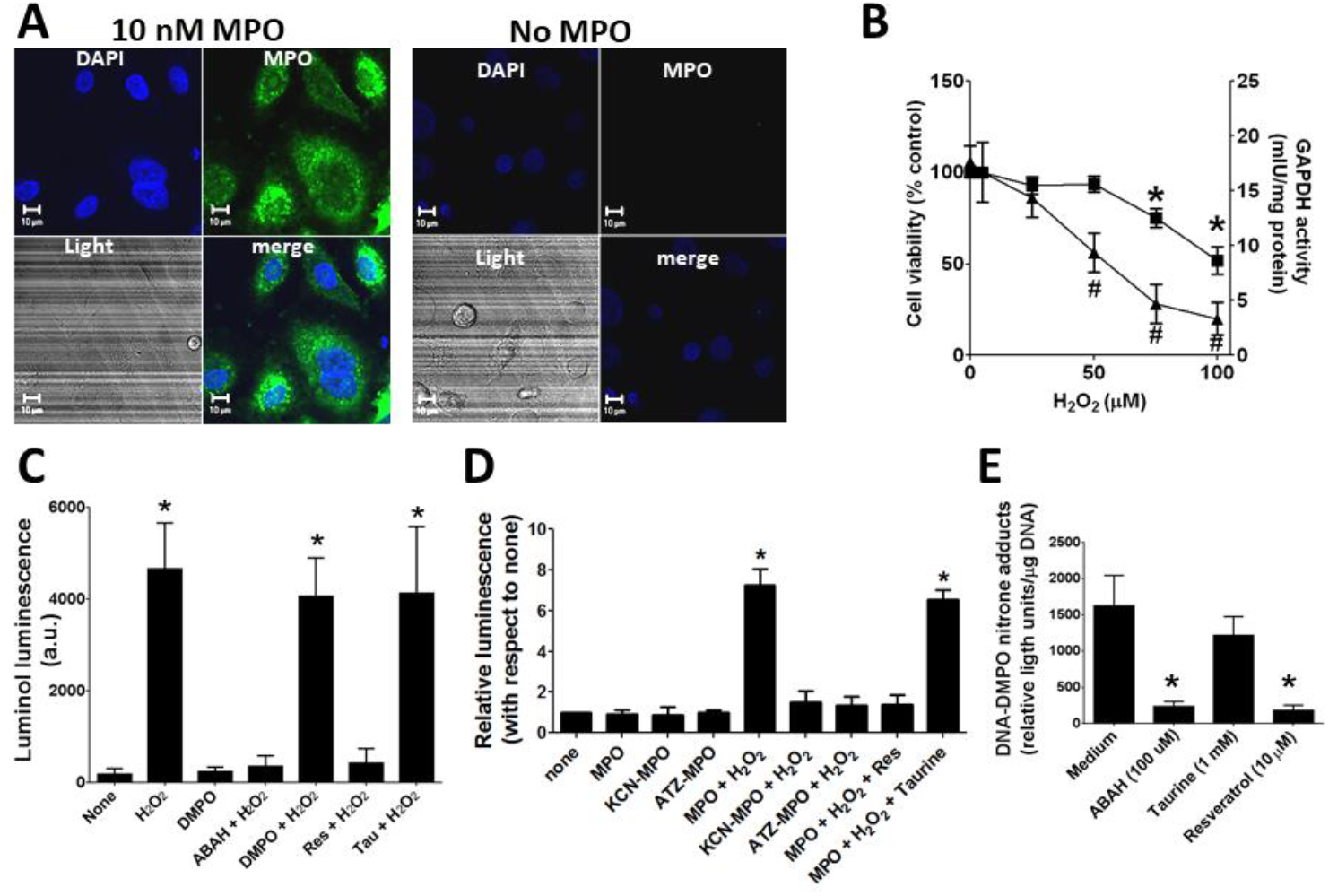
Uptake of MPO and intracellular production of HOCl in A549 epithelial cells. **A**) Single planar confocal image of MPO in A549 cells loaded with human MPO. **B**) Effect of intracellularly produced HOCl on cell viability and GAPDH activity. Cells were loaded with MPO and incubated with different concentrations of H_2_O_2_ for 30 min in HBSS^+^ containing 140 mM NaCl. After 30 minutes, the cells were rinsed to eliminate excess H_2_O_2,_ and cell viability was assessed using the MTT assay (closed squares). GAPDH activity, a key enzyme involved in the glycolytic pathway, was determined using a commercial kit in cell homogenates (closed triangles). **C**) Effect of MPO inhibition or scavenging of HOCl on HOCl-induced oxidation of luminol. Experiments were performed in cells loaded with MPO and then treated or not (none) with H_2_O_2_ in the presence or absence of 50 mM DMPO, 100 μM ABAH, 10 μM resveratrol (Res), and 1 mM taurine. Intracellular production of HOCl inside cells was triggered by the addition of 50 μM H_2_O_2_ and was assessed using luminol as a probe. **D**) Effect of the inactivation of MPO or scavenging of HOCl on luminol oxidation by intracellularly produced HOCl. Cells were loaded with either native or inactivated MPO and then treated or not with 50 μM H_2_O_2,_ and luminol oxidation was assessed by chemiluminescence. **E)** Effect of MPO inhibition or scavenging of HOCl on DNA-DMPO adduct formation. Cells loaded with active MPO were incubated in medium containing either 100 μM ABAH, 1 mM taurine, or 10 µM resveratrol. Then, HOCl production was triggered by a bolus addition of 50 μM H_2_O_2_. DNA-DMPO nitrone adducts were measured by ELISA and referred to DNA bound to the culture plate. Asterisks indicate p<0.05 with respect to none (no MPO, no H_2_O_2_). Data are representative or mean values ± s.e.m. from 4 independent experiments. a.u., arbitrary units.

To corroborate the production of HOCl inside the cells loaded with MPO and treated with H_2_O_2,_ we used luminol, a cell-permeable fluorescent probe that reacts with reactive biochemical species; however, the luminescence caused by reaction with HOCl is more stable than that generated, for example, with peroxynitrite (**Fig. 3C**). To support these data, we tested whether resveratrol or taurine can block the reaction with the probe. Taurine is an amine that reacts with HOCl but does not cross cell membranes; however, resveratrol is a *trans*-stilbene that crosses membranes and reacts quickly with HOCl before it reacts with luminol [7]. We observed that 1 mM taurine does not affect the luminescence signal, whereas 10 μM resveratrol completely blocks the reaction of HOCl with the probe inside the cell (**Fig. 3C**).

We then measured the production of HOCl inside A549 cells loaded with native and inactivated MPO after treatment with 50 μM H_2_O_2_. We observed that native and active MPO is needed to increase luminescence and that it was inhibited by resveratrol but not by taurine (**Fig. 3D**). Likewise, luminescence and nitrone adduct formation were inhibited when ABAH or resveratrol, but not taurine, was added to the reaction medium (**Fig. 3E**).

Previously, using a Fenton-like system (Cu^2+/^H_2_O_2_) to radicalize DNA, we found that DMPO prevents 8-oxo-dGuo formation [13]. Similar experiments in HOCl-induced DNA-centered radicals have not yet been reported. Herein, we used the G/GO system to produce a continuous flow of H_2_O_2_ to outcompete cell antioxidants (e.g., L-ascorbate and reduced glutathione) (**Fig. 4A**). These experiments were performed in the presence or absence of DMPO to study the production of 8-oxo-dGuo and DNA-DMPO nitrone adducts, respectively. The production of H_2_O_2_ in cells loaded with MPO and in the absence of DMPO resulted in 8-oxo-dGuo production, whereas in the presence of DMPO, DNA-centered radicals were trapped, and no 8-oxo-dGuo was produced (**Fig. 4A**). To produce DNA-DMPO nitrone adducts in MPO-loaded A549 epithelial cells, H_2_O_2_ and DMPO were needed. We found that active MPO, Cl^−^ and H_2_O_2_ were needed to produce 8-oxo-dGuo and DNA-DMPO nitrone adducts in the absence and presence of DMPO, respectively (**Fig. 4B**). Inhibition of MPO, with KCN or ATZ, before loading of A549 epithelial cells did not cause 8-oxo-dG or DNA-DMPO nitrone adducts.

**Figure 4:**
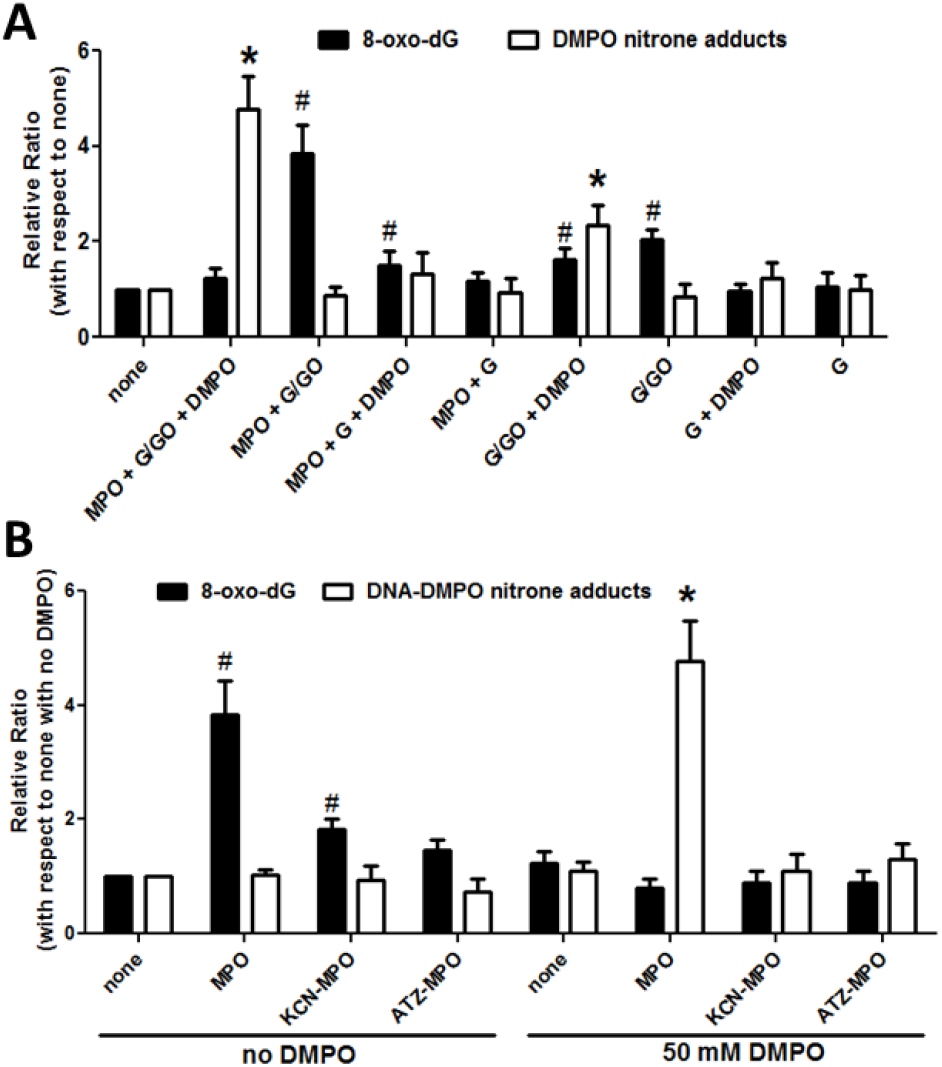
DMPO prevents the accumulation of 8-oxo-dGuo induced by intracellularly produced HOCl. **A**) Quantification of 8-oxo-dGuo and DMPO-nitrone adducts in DNA isolated from A549 cells in which HOCl was intracellularly generated in the absence or presence of DMPO. Intracellular generation of HOCl was triggered by a continuous flow of H_2_O_2_ by the glucose/glucose oxidase (5.6 mM G/1 mIU mL^-1^ GO) system. Reactions were stopped by rinsing the cells with 100 μM resveratrol in Tris-buffered saline, pH 7.8. **B**) Same as A, but cells were loaded with native MPO or MPO that was previously inactivated with KCN or ATZ. Reactions were performed in either the presence or absence of 50 mM DMPO to measure DNA-DMPO nitrone adducts or 8-oxo-dGuo, respectively. Data are shown as the mean values ± s.e.m. from three independent experiments. Asterisk (*) indicates a difference in DNA-DMPO nitrone adducts with respect to none. The pound symbol (#) indicates a difference in 8-oxo-dGuo with respect to none.

Genomic damage is sensed by the genome guardian p53, a transcription factor that controls the expression of genes involved in cell cycle arrest [31]. In cells loaded with MPO and treated with H_2_O_2_, p53 accumulated in the cell nucleus (**Fig. 5**), suggesting that the genomic damage induced by intracellularly produced HOCl was sensed.

**Figure 5.**
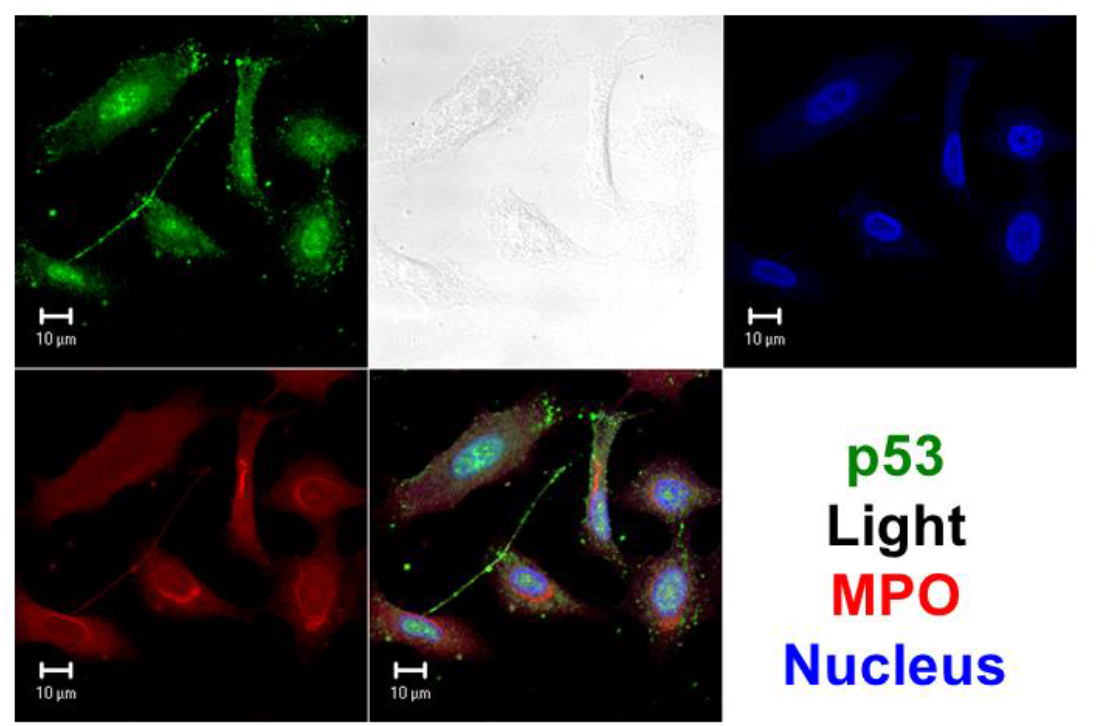
Intracellular generation of HOCl leads to the nuclear accumulation of p53. Planar confocal image showing the accumulation of p53 in A549 lung epithelial cells loaded with MPO and then treated with the 5.6 mM G/1 mIU/mL GO system for 15 min. A representative image of three separate experiments is shown.

**Figure 6:**
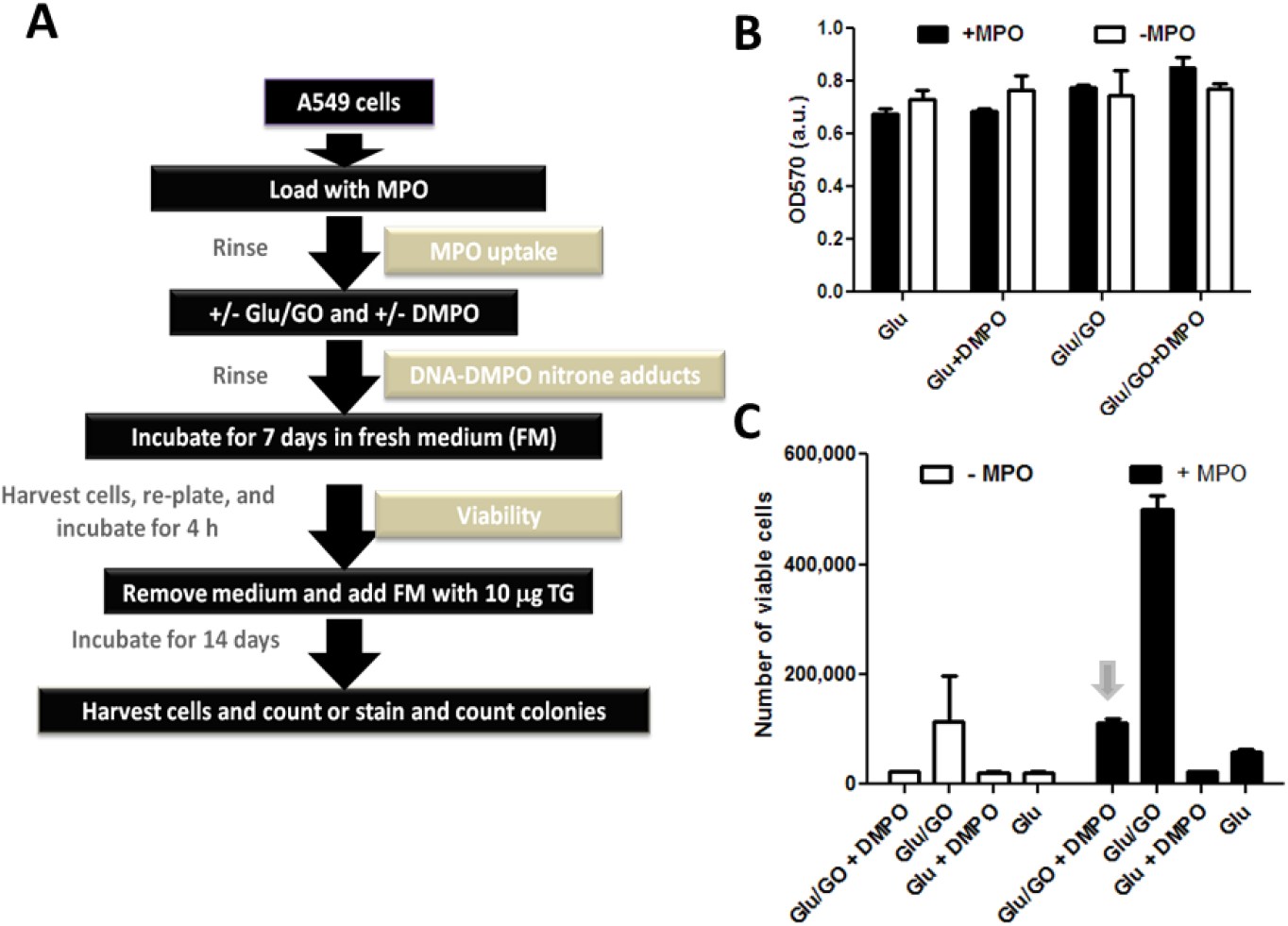
DMPO prevents HOCl-induced mutation of the *hrpt* gene in A549 epithelial cells. **A**) Scheme of the mutagenesis assay from the loading of A549 cells with MPO until the 6-TG selection assay (see Methods). **B**) MTT viability of A549 epithelial cells after 7 days of culture. These cells were treated as follows: loaded with 10 nM MPO, exposed to a flow of H_2_O_2_ generated by the 5.6 mM G/1 mIU/mL GO biochemical system with or without 50 mM DMPO for 15 min and then incubated for 7 days in fresh medium. **C**) Trypan blue exclusion assay for measurement of 6-TG-resistant cells (mutant cells) after 14 days of culture. Data show mean values ± s.e.m. gathered from three independent experiments. The arrow in the C panel indicates the effect of DMPO on the 6-TG-resistant cell count.

The *hrpt* gene is one of the most sensitive genes to the damage caused by mutagens. **Figure 6A** shows the workflow used to detect *hrpt* gene mutagenesis using the 6-TG reagent to select mutant cells. None of the cell treatments, loaded or not with MPO and with or without DMPO, caused significant cell death compared with cells treated with glucose only (**Fig. 6B**). After determination of cell viability, the medium was removed and replaced by fresh medium containing 10 μM 6-TG and incubated for 14 days to select *hrpt* mutant cells. After this incubation, the cells were carefully harvested, and a trypan blue exclusion assay was performed to distinguish the death (6-TG-sensitive, nonmutant cells) of living cells (6-TG-resistant, mutant cells) (see **Fig. 6A**).

These experiments showed that treating MPO-loaded A549 cells with a continuous flow of H_2_O_2_ resulted in the greatest generation of 6-TG-resistant cells. Mutagenesis, as well as the generation of 8-oxo-dGuo, was prevented when DMPO was present within the cell when DNA radical formation occurred (**Fig. 6C**).

## DISCUSSION

Herein, we used an *in vitro* experimental model to study the role of DNA radical formation at sites of neutrophilic inflammation in mutagenesis. Our data are consistent with the critical role of DNA radicalization in HOCl-induced mutagenesis in epithelial cells. Indeed, if not trapped by DMPO, DNA-centered radicals decay, amongst other DNA oxidation products, to 8-oxo-dGuo causing mutations, for example in the *hrpt* gene.

Upon pathogen infection or irritant exposure, local macrophages and other cells sense the insult and produce a wide panel of inflammatory mediators, such as cytokines and chemokines, that stimulate the nearby microvasculature and attract a large number of neutrophils to migrate across the vascular wall and infiltrate into tissues [32]. Neutrophilic inflammation [1] is an important acute response to irritation whose main role is to eliminate infection and enhance the adaptive immune response and tissue repair [33]. However, the role of HOCl produced by MPO in carcinogenesis has been proven *in vitro* and in experimental models [17]. Moreover, those patients who express more MPO due to a single nucleotide polymorphism in the promoter region (-463 G>A, rs#2333227), the GG genotype of the enzyme that is the most frequent in humans (approximately 45-60%), are the most susceptible to lung cancer [34], as well as other chronic inflammatory diseases in which MPO plays a pathogenic role [35]. The recent SARS-CoV-2 pandemic highlighted the importance of neutrophilic inflammation in genomic damage in airway cells [28]. Consequently, the study of the mechanism linking neutrophilic inflammation-induced DNA damage is of fundamental importance to reduce the potential genotoxicity and mutagenicity in the lungs of COVID-19 patients.

MPO is the main enzyme contained in azurophilic granules of neutrophils that are released upon activation [36]. MPO can be taken up by cells in inflamed tissue, where HOCl production can damage the genome (**Fig. 7**). MPO is the only animal peroxidase that, under physiological concentrations of halides and pH, produces HOCl by H_2_O_2_-driven oxidation of chloride anions [37, 38]. Depending on H_2_O_2_ concentrations, MPO operates either by a peroxidase or a chlorinating cycle at high or low H_2_O_2_ concentrations, respectively [39]. The peroxidase cycle of MPO is important in radical production, whereas the chlorinating cycle of MPO results in complex I oxidation of chloride anion into HOCl [40]. The chlorinating cycle of MPO is the most important at sites of neutrophilic inflammation and results in increased chlorination of proteins, lipids, and nucleic acids [1, 40]. Physiological concentrations of Cl^−^ were used in our work to show the production of HOCl in biochemical systems and A549 epithelial cells. Notably, the concentration of HOCl found at sites of neutrophilic inflammation has been estimated to be between 30 and 200 μM [41], which is higher than the concentrations we used in this study.

**Figure 7.**
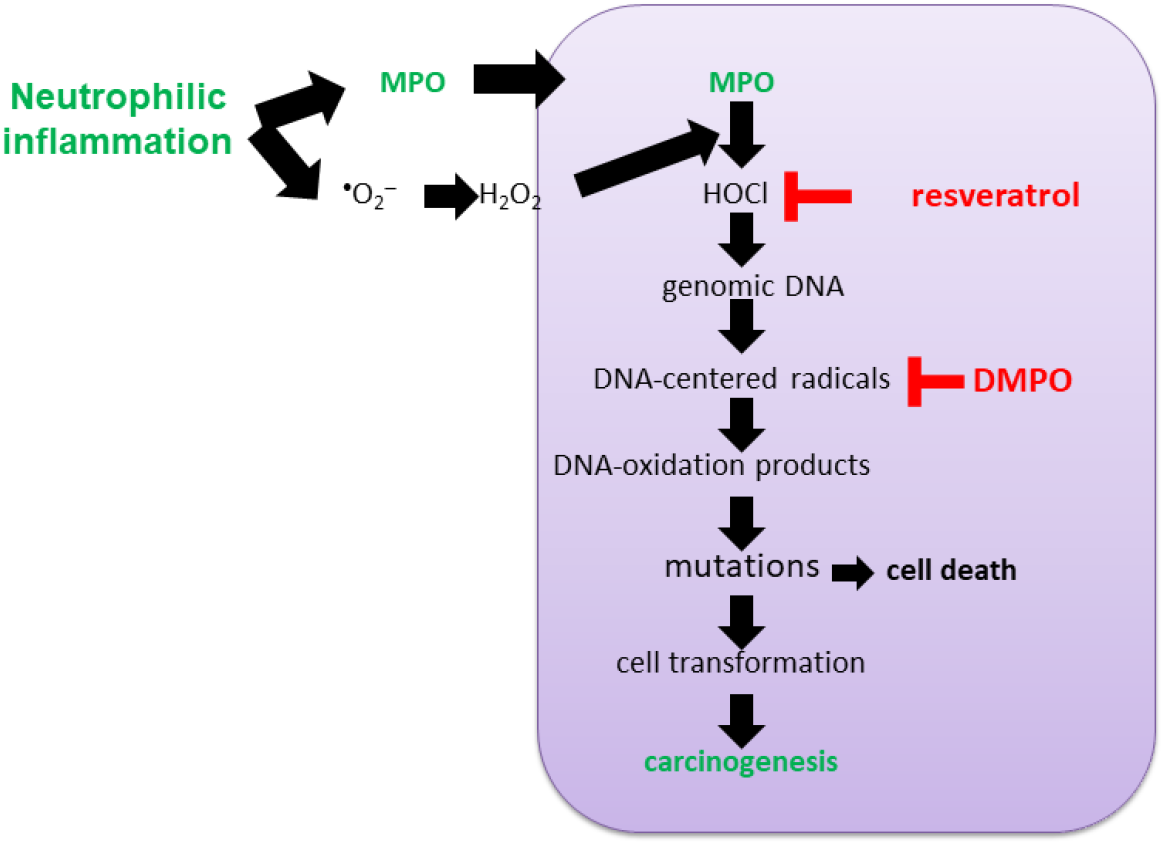
The potential impact of DNA radicalization caused by intracellularly produced HOCl in airway epithelial cells at sites of neutrophilic inflammation on mutagenesis. The *hrpt* gene mutation was due to intracellularly produced HOCl but not extracellularly produced HOCl. Our findings establish the critical role of the transient formation of DNA radicalization in the mutagenesis produced by HOCl generated inside epithelial cells, which may explain neutrophilic inflammation-induced carcinogenesis. Inhibitors of MPO (*e.g.,* ABAH), cell-permeable scavengers of HOCl (*e.g.,* resveratrol) or nitrone spin traps (*e.g.,* DMPO or its derivatives) can block mutagenesis in neutrophilic inflammation and thus prevent carcinogenesis in the lung exposed to chemical, physical or biological airborne irritants (reviewed in [8]).

Neutrophil activation, for example, during neutrophilic inflammation in the lung exposed to bacterial endotoxin, involves the activation of NADPH-oxidase-2 and further NET formation [42]. Damage caused by activated neutrophils in surrounding tissues involves oxidative modification and chlorination of proteins, lipids, and nucleic acids [35]. However, neutrophils are protected against HOCl toxicity because of their high intracellular concentration of taurine (20-50 mM), which reacts faster with HOCl than any other biological target [43].

The reaction of HOCl with DNA produces DNA chloramines that quickly decay, forming DNA radicals centered in nitrogen atoms [20]. The unpaired electron can be transferred to other atoms in the structure of each nucleotide to produce damage to the backbone, which results in DNA fragmentation, base chlorination (*e.g.,* Cl-dG and Cl-dA) and other oxidation products, such 8-oxo-dGuo [8, 11, 21]. Resveratrol is a *trans*-stilbene that easily crosses cell membranes and reacts rapidly with HOCl before it reacts with luminol or genomic DNA [7]. Indeed, the rate constant for the one-electron oxidation reaction of luminol by HOCl has been estimated as approximately 10^-7^ s^-1^ (*Personal communication, Dr Michael Ashby, Department of Chemistry and Biochemistry, The University of Oklahoma at Norman*). For the first time, we found that by trapping DNA radicals with DMPO, HOCl-induced 8-oxodGuo formation in cells was blocked. We found similar results in trapping protein-centered radicals with DMPO, observing that DMPO reduces HOCl-induced GAPDH fragmentation [9] and reduces carbonyls—one of the end products of free radical damage to proteins, in macrophages activated with lipopolysaccharide [44].

Upon genomic damage, p53 is translocated into the cell nuclei, where it binds to response elements located at regulatory sequences of genes involved in arresting the cell cycle until cell survival or death is decided [23]. If this mechanism fails, mutant cells can escape this vigilance mechanism, and subsequently, cell transformation and carcinogenicity may occur [24]. We found that HOCl produced by active MPO inside A549 epithelial cells results in p53 translocation inside the cell nuclei. These results suggest that oxidative damage caused by HOCl is sensed and that a proper response can be triggered.

Mutagenesis of the *hrpt* gene is one of the most sensitive genes to mutagenic insults [26]. To test whether HOCl triggers mutagenesis of the *hrpt* gene, a glucose/glucose oxidase system was used to generate a nontoxic but continuous flow of H_2_O_2_ to overwhelm antioxidant defense mechanisms [7]. We found that intracellularly generated HOCl increased the number of 6-TG-resistant cells. These mutant cells proliferate during the 14 days of incubation with 6-TG. These mutant cells, as well as the 8-oxo-dGuo content in their genomic DNA, were reduced if DNA-centered radicals were trapped with DMPO [8]. Our mutagenesis data are consistent with a nonmutagenic effect of DNA-DMPO nitrone adducts in the genome. We are still not sure whether, and if so, how, DNA-DMPO nitrone adducts can be repaired. However, Bhattacharjee *et al.* [45, 46] demonstrated using a model of macrophages treated with a hydroxyl radical-generating system that these DNA-DMPO nitrone adducts can be repaired over a short period, most likely through the base-excision repair pathway. They also found that DNA repair enzymes modify DNA damage, including the removal of adducted DMPO, and there are multiple and overlapping DNA repair pathways. This is great news because we can stop HOCl-induced mutagenesis, but it is bad news if our interest is to detect DNA-DMPO nitrone adducts because they may be underestimated [8].

Altogether, our findings establish the causative role of transiently formed DNA-centered radicals during the mutagenesis produced by HOCl generated inside epithelial cells, which may explain neutrophilic inflammation-induced carcinogenesis. Inhibitors of MPO, cell-permeable scavengers of HOCl (*e.g.,* resveratrol), or nitrone spin traps may help to prevent the mutagenesis process caused by intracellularly produced HOCl (**Fig. 7**).

## Abbreviations

8-oxo-dGuo: 8-oxo-7,8-dihydro-2’-deoxyguanosine
6-TG: 6-thioguanine
ABAH: 4-aminobenzoic acid hydrazide
ATZ: 3-amino-1,2,4-triazole
DMPO: 5,5-dimethyl-1-pyrroline-*N* oxide
G/GO: glucose/glucose oxidase
HBSS^-^: Hank’s buffered saline solution without Ca^2+^ and Mg^2+^
HBSS+: Hank’s balanced saline solution with Ca^2+^ and Mg^2+^
IST: immune-spin trapping
hrpt: hypoxanthine phosphorylbosyl transferase
MPO: myeloperoxidase
NET: neutrophil extracellular trap
NOX-2: NADPH oxidase-2
PMA: phorbol 12-myristate 13-acetate
SHA: salicylhydroxamic acid
res: resveratrol
tau: taurine

## ACKNOWLEDGMENTS

This research was supported in part by the Agencia Nacional para la Promoción de la Ciencia y la Tecnología, FONCYT, Argentina (PICT-2018-3435 and PICT-2021-I-A-000147 to DCR and SEGM), The Consejo Nacional de Investigaciones Científicas y Técnicas (CONICET), Argentina (PUE013 IMIBIO-SL) and the Universidad Nacional de San Luis, Argentina (PROICO 02-3418 to DCR and PORICO 10-0218 to SEGM).

